# The genus *Microniphargus* (Crustacea, Amphipoda): evidence for three lineages distributed across northwestern Europe and transfer to Pseudoniphargidae

**DOI:** 10.1101/2020.08.25.266817

**Authors:** Dieter Weber, Fabio Stoch, Lee R.F.D. Knight, Claire Chauveau, Jean-François Flot

## Abstract

*Microniphargus leruthi* Schellenberg, 1934 (Amphipoda: Niphargidae) was first described based on samples collected in Belgium and placed in a monotypic genus within the family Niphargidae. However, some details of its morphology as well as recent phylogenetic studies suggest that *Microniphargus* may be more closely related to *Pseudoniphargus* (Amphipoda: Pseudoniphargidae) than to *Niphargus.* Moreover, *M. leruthi* ranges over 1,469km from Ireland to Germany, which is striking since only a few niphargids have confirmed ranges in excess of 200km. To find out the phylogenetic position of *M. leruthi* and check whether it may be a complex of cryptic species, we collected material from Ireland, England and Belgium then sequenced fragments of the mitochondrial cytochrome *c* oxidase subunit 1 gene as well as of the nuclear 28S ribosomal gene. Phylogenetic analyses of both markers confirm that *Microniphargus* is closer to *Pseudoniphargus* than to *Niphargus*, leading us to reallocate *Microniphargus* to Pseudoniphargidae. We also identify three congruent mito-nuclear lineages present respectively in Ireland, in both Belgium and England, and in England only (with the latter found in sympatry at one location), suggesting that *M. leruthi* is a complex of at least three species with a putative centre of origin in England.

## Introduction

*Microniphargus leruthi* Schellenberg, 1934 (family Niphargidae) was first described from Engihoul Cave in Wallonia (Belgium) and placed into a new, monotypic genus considered as closely related to the genus *Niphargus* Schiødte, 1849 (Schellenberg 1934). *M. leruthi* is characterised by its small body size (1.2-1.5mm in length), the scant setulation of its mandibular palps, an evident protrusion on the carpus of its gnathopods (particularly pronounced on the first pair of gnathopods: Knight & Gledhill 2010) and its telson widely incised with an angle of around 80° in its indentation, all of which were used to justify the erection of a new genus (Schellenberg 1934). However, several other genera similarly erected on the basis of distinctive morphological features of unknown variability have been synonymized with *Niphargus* when molecular data became available (see e.g. Borko *et al.* 2020).

Although *M. leruthi* is presently placed in the family Niphargidae, the shape of its telson is quite similar to that of the genus *Pseudoniphargus* Chevreux, 1901, which is placed in a different family (Pseudoniphargidae) together with the monotypic *Parapseudoniphargus* Notenboom, 1988. The taxonomic position of the family Pseudoniphargidae, defined on vague morphological characters, has long been controversial, having been included in the superfamilies Hadzioidea, Niphargoidea, Crangonyctoidea, Gammaroidea or included in the families Gammaridae and Melitidae (see Notenboom 1988a for a detailed analysis). Notenboom (1988b) in his cladistic analysis placed the family within the families Eriopisidae and Melitidae, whereas Allocrangonyctidae (comprising two stygobitic species from North America) were later considered as the most closely related family. In fact, in their recent revisions of Amphipod taxonomy, Lowry & Myers (2012, 2017) included the family Pseudoniphargidae within the superfamily Allocrangonyctoidea, while Niphargidae were allocated to Crangonyctoidea. Recent molecular studies have rejected this hypothesis, suggesting that Pseudoniphargidae are the sister group of Niphargidae (Jurado-Rivera et al. 2017, Moškrič & Verovnik 2019, Copilas-Ciocianu et al. 2020). Although numerous mitochondrial sequences of *Pseudoniphargus* are available, there are only three partial 28S sequences for this genus and no genetic data at all for the family Allocrangonyctidae and for the genus *Parapseudoniphargus*, hindering a definitive taxonomic assessment of this clade.

Existing molecular data regarding *M. leruthi* are also scarce, with only 10 sequences available in GenBank so far (Fišer *et al.* 2017; Moškrič & Verovnik 2019), all of which from nuclear markers. No mitochondrial sequence for specimens of *Microniphargus* has been published so far. Moškrič & Verovnik (2019) recovered a (*Microniphargus* + *Pseudoniphargus*) clade as a sister group to *Niphargus* using one protein-coding nuclear gene; however, another protein-coding nuclear marker in the same study yielded a discordant position of *Microniphargus* within *Niphargus.* More recently Copilaş-Ciocianu *et al.* (2020), in a large-scale phylogeny of amphipods comprising one *Microniphargus*, two *Pseudoniphargus* and two *Niphargus* species, also recovered *Microniphargus* as more closely related to *Pseudoniphargus* than to *Niphargus.*

*Pseudoniphargus* comprises 71 stygobitic species (Stokkan et al. 2018), all strict endemics present in North Africa and Benin, the Mediterranean region, the Iberian Peninsula, the archipelagos of Canaries, Madeira and Azores, and two species in Bermuda, whereas *Parapseudoniphargus* comprises a single, stygobitic species from southern Spain. By contrast, *Microniphargus leruthi* is found in north-western Europe: Belgium (Leruth 1939; Spangenberg 1973; Karaman & Ruffo 1986; Delhez *et al.* 1999), Germany (Spangenberg 1973; Karaman & Ruffo 1986; Fuchs 2007; Matzke *et al.* 2009; Stein *et al.* 2012), Luxembourg (Hoffmann 1963) as well as Ireland (Arnscheidt *et al.* 2008; Knight & Penk 2010; Knight & Gledhill 2010) and Great Britain (Knight & Gledhill 2010). The very large range of *M. leruthi* (over 1,469 km) is unusual as only a few niphargids have ranges exceeding 200km (Trontelj *et al.* 2009): some species previously considered to be wide-ranging, such as *Niphargus aquilex* Schiödte, 1855 and *Niphargus virei* Chevreux, 1896, have been found to be complexes of cryptic species (McInerney et al. 2014; Lefébure et al. 2006). The only species with confirmed ranges more extended than *M. leruthi* are *Niphargus hrabei* S. Karaman, 1932 (>1,300km) and *Niphargus valachicus* Dobreanu & Manolache 1933 (>3,200km), two epigean species with enhanced dispersal via surface water (Copilaş-Ciocianu *et al.* 2017). The wide range of *M. leruthi* could therefore be due to the presence of undetected species boundaries.

To resolve these uncertainties, we conducted a molecular study on *M. leruthi* collected in Ireland, England and Belgium using both 28S (nuclear) and COI (mitochondrial) markers. Our aims were (i) to confirm the phylogenetic position of *Microniphargus* relative to the genera *Niphargus* and *Pseudoniphargus* and (ii) to test for the possible existence of cryptic lineages within *M. leruthi*.

## Material and methods

### Sampling and sequencing

Although we carried out intensive and targeted sampling for *M. leruthi*, especially in caves around the type locality, we were only able to collect it at a single location on the European continent: the Grotte de Comblain (Wallonia, Belgium), which is 20 km away from the type locality. We also collected *M. leruthi* from one site in Ireland (Polldubh in Galway) and two sites in England (Sweetwater Pot in South Devon and Swildon’s Hole in Somerset; Fig. 1). All the material (Table 1) was determined morphologically by one of us (L. K.). Specimens were collected by sweeping a long-handle net fitted with a 250 μm mesh collecting bag along the bottom and sides of cave pools, making sure to disturb the substrate in order to suspend both sediment and specimens into the water column. The collected specimens were immediately preserved in 96% ethanol and kept at −20°C until DNA was isolated.

**Figure 1:**
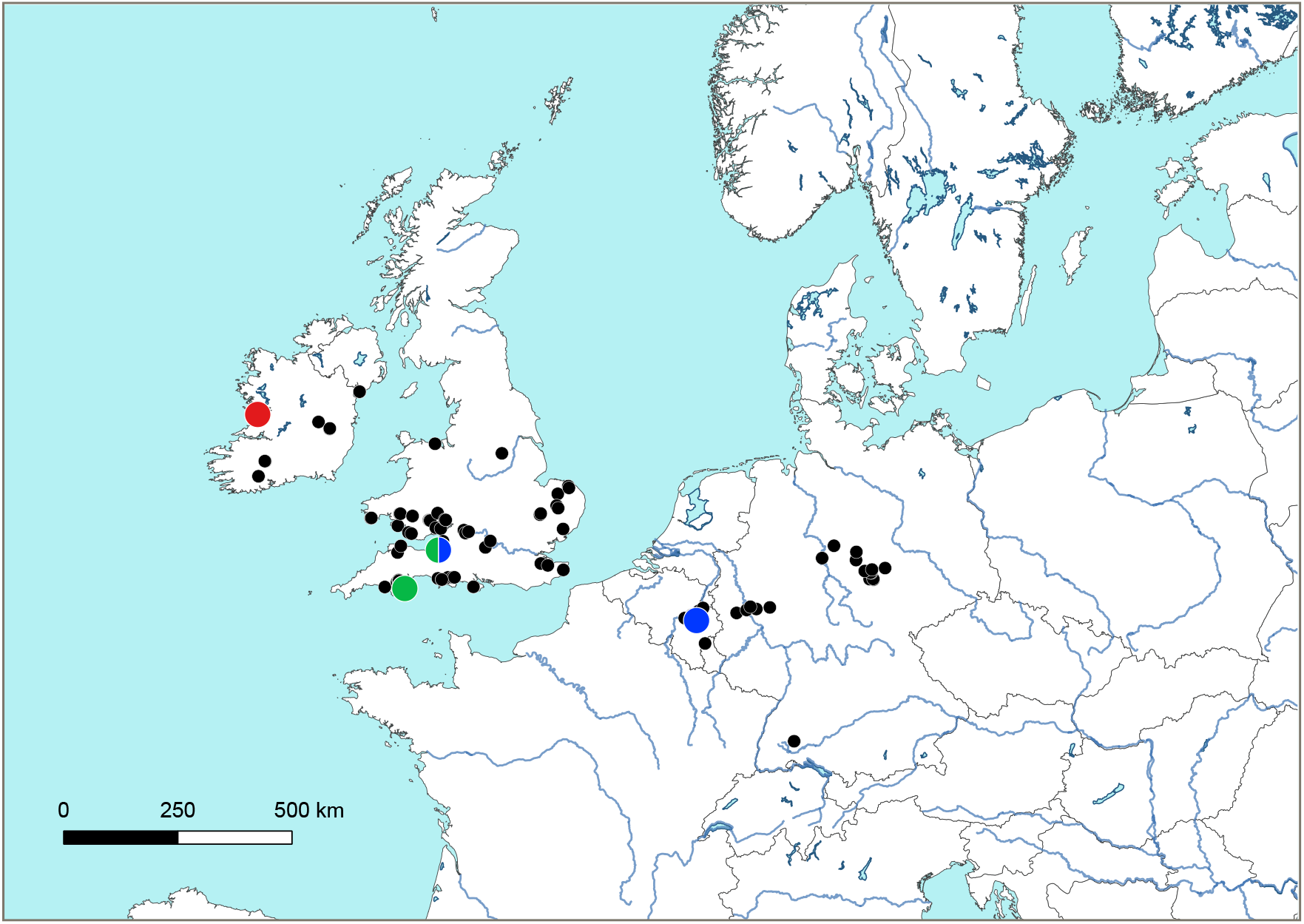
Map showing the distribution of *Microniphargus leruthi*. Black dots: literature data based on morphological determination (including https://hcrs.brc.ac.uk/hcrs-database, accessed 27th May 2020). Red: lineage A; blue: lineage B; green: lineage C.

**Table 1:**
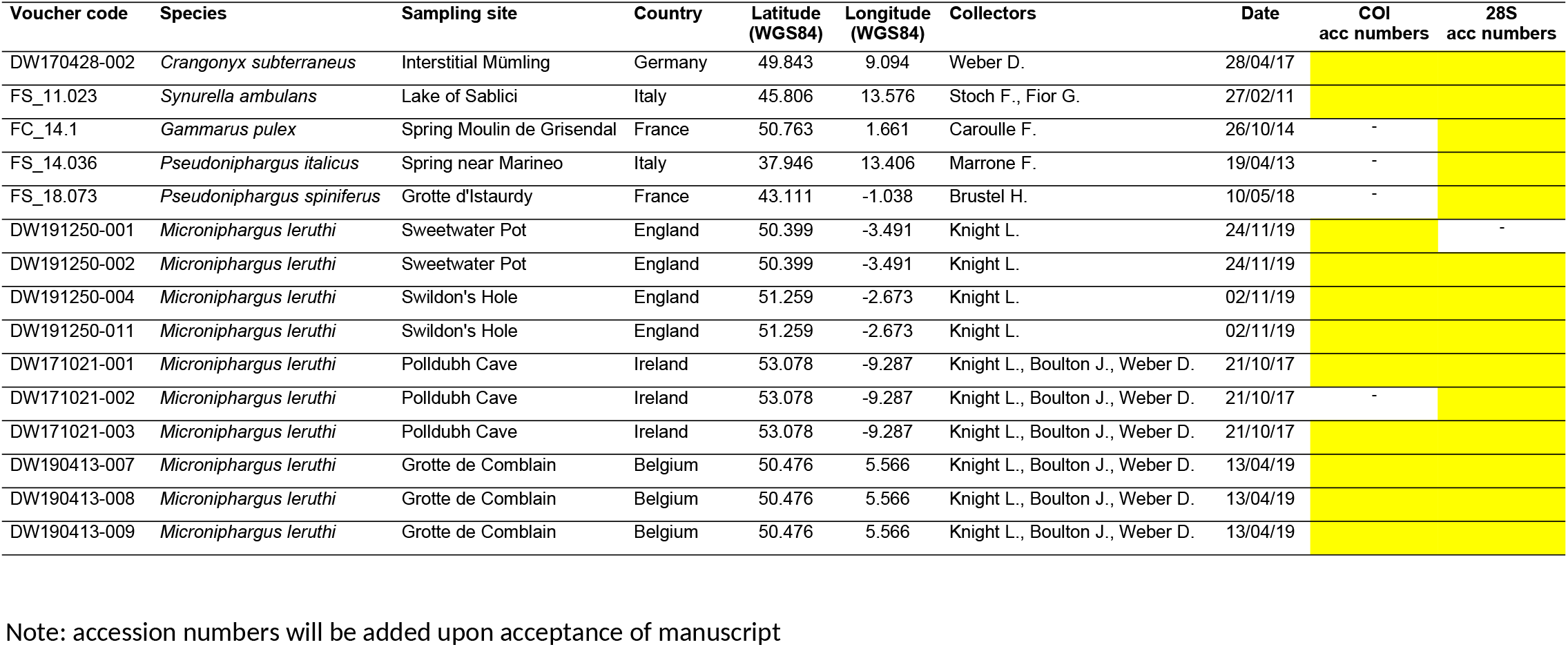
Material newly sequenced for the present study

Due to the small size of *M. leruthi*, we used one whole specimen for each DNA isolation. DNA was extracted following the standard protocol of the NucleoSpin® Tissue Kit (Macherey-Nagel) except that we performed two elution steps, the first one with 60 μL and the second with 40 μL (instead of a single elution step with 100 μL) to achieve a higher concentration of DNA. The resulting DNA isolates are stored at −20°C in the collections of the Evolutionary Biology & Ecology research unit at the Université libre de Bruxelles (ULB).

The Folmer fragment of the cytochrome *c* oxidase subunit 1 (COI) gene was amplified via polymerase chain reaction (PCR) (Folmer *et al.* 1994) using the primers HCO2198-JJ and LCO1490-JJ (Astrin & Stüben 2008; see Table 2). The PCR mix contained 1XL DNA template (variable concentration), 0.8XL of each primer (10pmol/XL), 5XL of DreamTaq DNA Polymerase (Thermo Scientific) and 2.4μL ultrapure water. PCR cycling conditions were an initial 3-min denaturation step at 94°C followed by 36 cycles of 20s denaturation at 94°C, 45s annealing at 50°C, and 60s extension at 65°C; then a final 2min elongation step at 65°C.

**Table 2:**
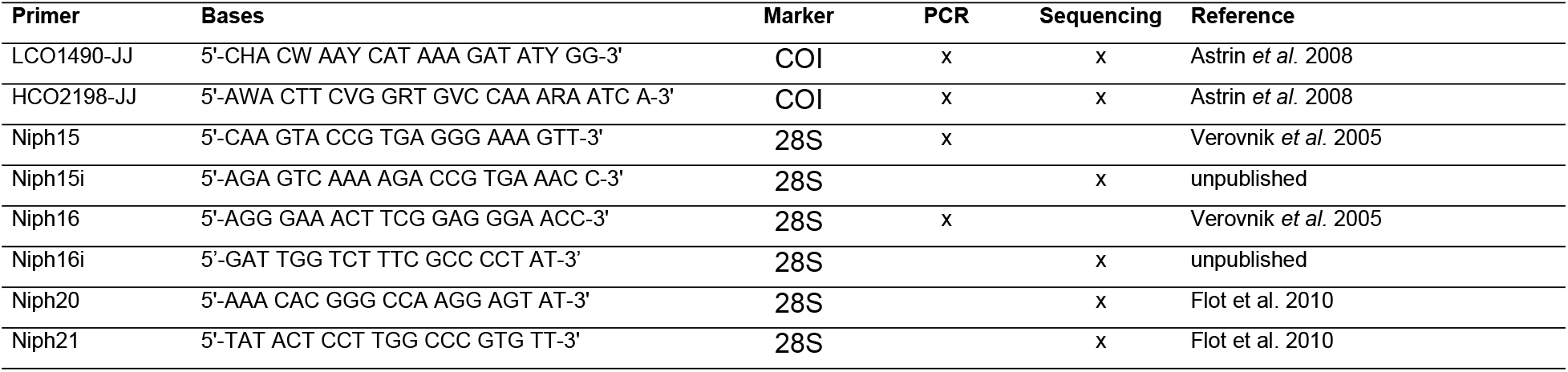
Primers used in the present study

We also sequenced Verovnik’s fragment of the nuclear 28S ribosomal gene. The primers Niph15 and Niph16 (see Table 2) were used for amplification (Verovnik *et al.* 2005). The PCR mix for 28S contained 2XL of DNA template (variable concentration), 1Xl of each primer (10pmol/XL), 0.2XL of REDTaq DNA Polymerase (Sigma-Aldrich), 5μL REDTaq reaction buffer and 15.8μL ultrapure water. PCR cycling conditions for 28S were an initial 3min denaturation step at 95°C; followed by 56 cycles of 30s denaturation at 94°C, 60 s annealing at 45°C, and 90s extension at 72°C.

The amplification success of each PCR reaction was verified using agarose gel electrophoresis, then PCR products were sequenced at Genoscreen (Lille, France). For COI the primer used for sequencing were the same as for PCR amplification, whereas for 28S we used the primers Niph20 and Niph21 (Flot, Wörheide, *et al.* 2010) as well as one or both of two new internal primers located slightly inward of the primers used for initial amplification (Niph15i and Niph16i; see Table 2).

The resulting chromatograms were assembled and cleaned using Sequencher version 4.1.4 (Gene Codes, USA). Whenever double peaks were observed in both the forward and reverse chromatograms of an individual, we considered this individual as polymorphic and called its two haplotypes “a” and “b” in downstream analyses.

### Phylogenetic analyses

We compiled comprehensive sets of COI and 28S including all sequences available in GenBank to date, then curated them manually to remove duplicates. The resulting set of 1384 COI sequences was aligned manually, whereas for the 255 sequences of 28S (including two gammarids *Gammarus fossarum* and *Gammarus pulex* and two crangonyctids *Crangonyx subterraneus* and *Synurella ambulans* as outgroups) we used MAFFT 7’s E-INS-i mode (Katoh *et al.* 2019).

The comprehensive 28S alignment was used to reconstruct a global phylogeny of niphargid and pseudoniphargid amphipods. The best-fit substitution model, selected using ModelFinder (Kalyaanamoorthy et al. 2017) according to the Bayesian Information Criterion (Schwarz 1978), was GTR+F+I+G4 (codes follow the IQTREE manual). Phylogenetic relationships were reconstructed using maximum likelihood with 1,000 ultrafast bootstrap replicates (Hoang et al. 2018) in IQ-TREE 2 (Minh et al. 2020); 253 out of 255 sequences (including all *Microniphargus* sequences) passed the gap/ambiguity test in IQTree 2 and were used in the analysis.

The comprehensive COI alignment was analysed using ABGD (Automatic Barcode Gap Discovery, available online at https://bioinfo.mnhn.fr/abi/public/abgd/), a distance-based species delimitation tool (Puillandre et al. 2012) that first attempts to infer the most likely position of a barcode gap (‘initial partitioning’) before conducting a second round of splitting by recursively applying the same procedure on the groups defined during the first step (‘recursive partitioning’). ABGD was run on the public webserver with default parameters.

A subset of the COI sequences (comprising all new *Microniphargus* sequences, all high-quality, complete *Pseudoniphargus* COI sequences inferred from complete mitochondrial genome sequences from Bauzà-Ribot et al. (2012) and Stokkan et al. (2016, 2018) plus two sequences of *Niphargus* and sequences of the Crangonyctidae *Crangonyx subterraneus* and *Synurella ambulans* (as outgroups) was used to build a ML tree using IQ-TREE 2 with the same modalities illustrated for 28S; the best-fit substitution model, selected using ModelFinder (according to the Bayesian Information Criterion was TIM+F+I+G4 (codes follow the IQTREE manual).

Phylogenetic networks were built for the COI and 28S sequences obtained from *Microniphargus* using HaplowebMaker (Spöri & Flot 2020, available online at https://eeg-ebe.github.io/HaplowebMaker/). Average genetic distances between *Microniphargus* sequences identified as belonging to different lineages were computed in MEGA X (Kumar et al. 2018) using p-distances.

## Results

For both COI and 28S, we successfully sequenced nine *Microniphargus leruthi* specimens. One *M. leruthi* individual (from Belgium) displayed a double peak in its 28S chromatograms and was therefore represented by two sequences in all downstream analyses, whereas other individuals were homozygous for the 28S marker. For COI, four individuals (one from Belgium and two from England) displayed a single double peak each and were therefore represented by two sequences “a” and “b” in all downstream analyses (Fig. 2).

**Figure 2:**
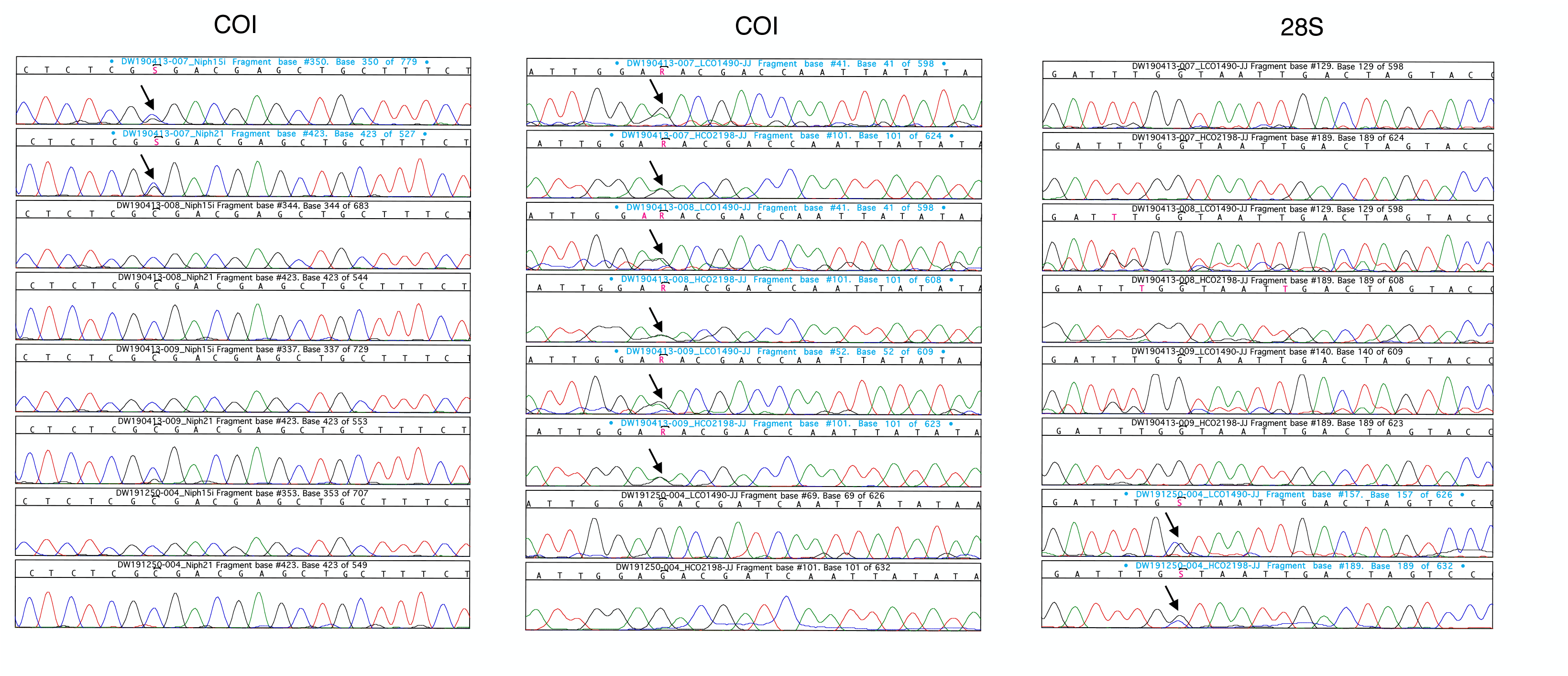
Screenshots of the Sequencher program showing the double peaks identified in the COI (left and middle panel) and 28S (right panel) chromatograms.

Our comprehensive 28S phylogeny including all published sequences of *Microniphargus*, *Niphargus*, *Pseudoniphargus* supported a (*Microniphargus* + *Pseudoniphargus*) clade with 100% of the ultrafast bootstrap replicates (Fig. 3). Our COI phylogeny confirmed this sister-clade relationship between *Pseudoniphargus* and *Microniphargus* (supported by 96% of the ultrafast bootstrap replicates) and revealed the latter to be composed of three main clades A (found only in Ireland), B (found both in Belgium and in England) and C (found only in England), with >99% ultrafast bootstrap support for each of them. Clade B comprised two subclades comprising respectively Belgian and English sequences, also with >99% ultrafast bootstrap support. Clade B and C co-occurred at one sampling site (Fig. 1).

**Figure 3:**
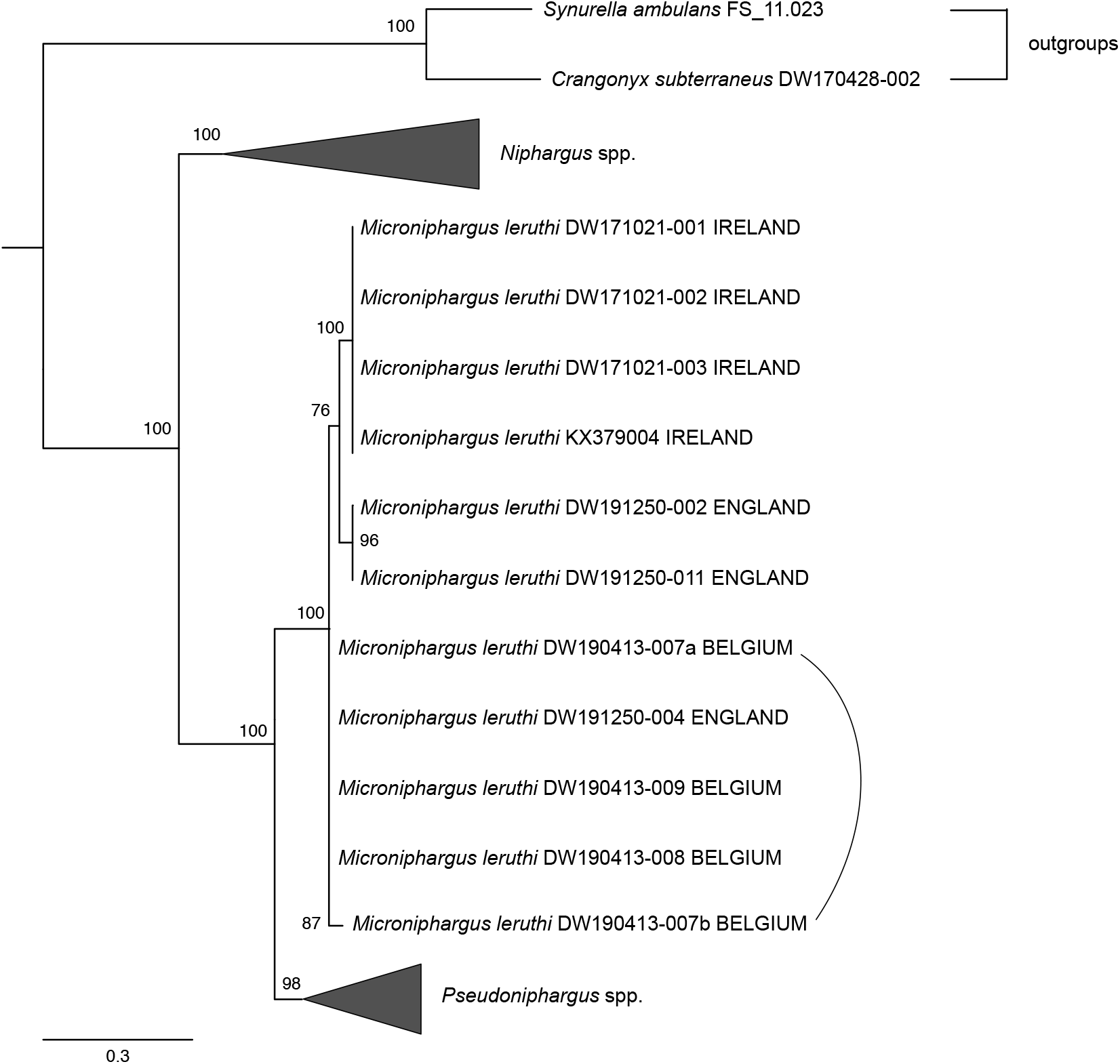
28S maximum-likelihood phylogeny of *Microniphargus*, *Niphargus* and *Pseudoniphargus* (with two crangonyctids as outgroups). The tree was turned into a haploweb by adding connections between haplotypes found co-occurring in the same individual.

The COI lineages A, B and C were separated by averages p-distances of 0.073: 0.081 between A and B, 0.072 between A and C, and 0.066 between B and C; whereas the p-distance between the two sub-lineages of C was 0.029. ABGD’s initial partitioning of our comprehensive COI dataset supported a three-species hypothesis for *Microniphargus leruthi*, whereas the recursive partitioning favoured a four-species hypothesis separating the Belgian and English sub-lineage of lineage C. The 28S haploweb revealed three fields for recombination (FFRs *sensu* Doyle 1995), i.e. putative species following the criterion of mutual allelic exclusivity (Flot et al. 2010) corresponding to clades A, B and C (Fig. 5).

**Figure 4:**
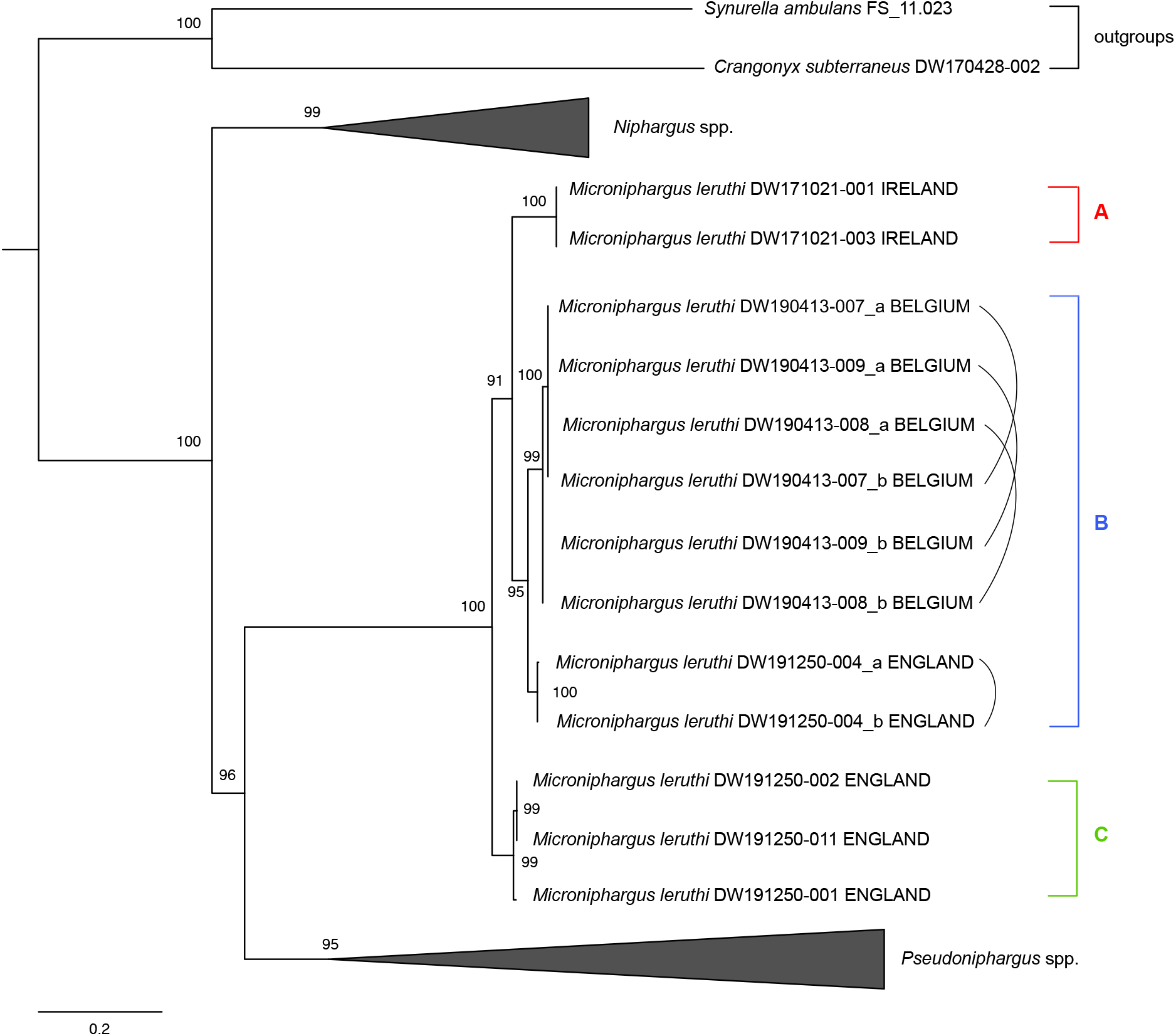
COI maximum-likelihood phylogeny of *Microniphargus* and *Pseudoniphargus* (with two *Niphargus* and two crangonyctids as outgroups). The tree was turned into a haploweb by adding connections between haplotypes found co-occurring in the same individual.

**Figure 5:**
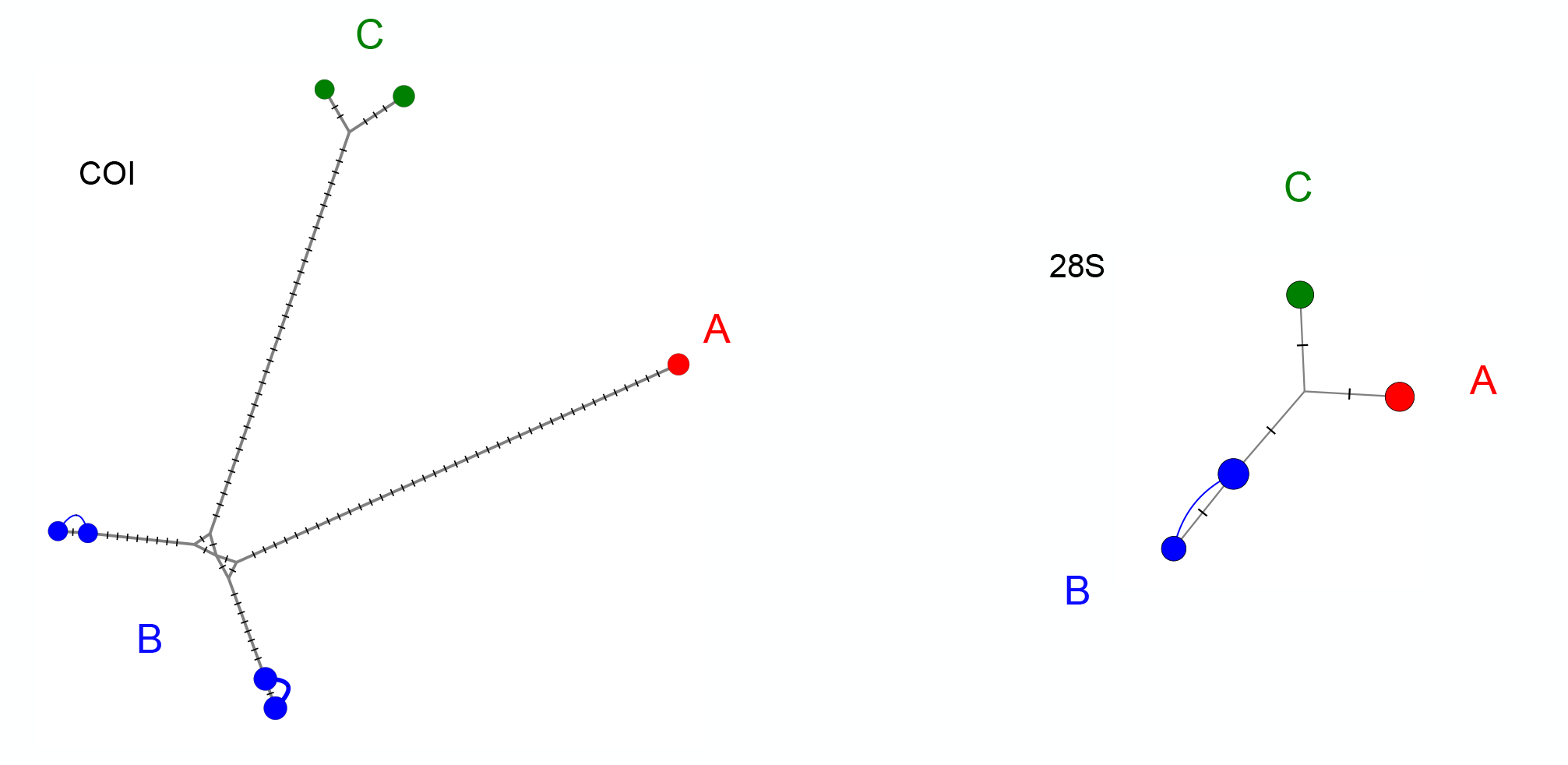
Median-joining networks of the *Microniphargus* COI and 28S sequences obtained in the present study. The networks were turned into haplowebs by adding connections between haplotypes found co-occurring in the same individual.

## Discussion

### Key novel, high-quality sequences were acquired

Our newly collected sequences include the first COI sequences of *Microniphargus leruthi* (and of *Crangonyx subterraneus*) made available to date, as well as new 28S sequences that significantly improve the currently available sequences for these two species: the single Verovnik’s 28S fragment sequence available till now for *C. subterraneus* (EU693288; Fišer *et al.* 2008) is 100% identical to ours (except for one obvious error at position 25), but its last 140 bp are lacking; the single 28S sequence of *M. leruthi* previously published (KX379004.1; Fišer *et al.* 2017) is 100% identical to our complete sequences from Ireland, but with the first 59 bp and last 156 bp lacking; whereas the three *Pseudoniphargus* sequences available till now were also highly incomplete. The high-quality 28S and COI sequences we obtained from representative individuals of *C. subterraneus* from Germany*, Pseudoniphargus italicus* from Sicily and *P. spiniferus* from Basses Pyrénées in France, as well as from each of the three lineages of *M. leruthi* identified in our study, will make it easier to include these species as outgroups in future studies of *Niphargus*, *Pseudoniphargus* and other related genera.

### Both COI and 28S sequences of *Microniphargus* were found to contain double peaks

Out of the nine *M. leruthi* individuals whose COI marker was sequenced, four (three from Belgium and one from England) presented a double peak in their COI chromatograms. For the three Belgian specimens it was a R = A or G transition in position 101, whereas for the English specimen it was S = C or G transversion in position 189. These mutations were not synonymous but corresponded to N (asparagine) ↔ D (aspartate) and A (alanine) ↔ G (glycine) mutations in the translation amino acid sequences. Such mitochondrial double peaks are rare in niphargids: for instance, no double peak was observed in the COI chromatograms of the hundreds of niphargids sequenced in Eme *et al.* (2018). The presence of two distinct COI sequences in *M. leruthi* individuals may be the result of heteroplasmy, i.e. the presence of two distinct mitochondrial lineages in the cells of an organism, or of a recent numt, i.e. a nuclear pseudogene of a mitochondrial sequence following the transfer and integration of a copy of this sequence in a nuclear chromosome (Dierckxsens *et al.* 2020). Determining which one of these two hypotheses is correct in the present case will require whole-genome sequencing, which is beyond the scope of the present study, but in any case the very limited divergence between the COI sequences found co-occurring in some individuals (with a single double peak per individual) did not hinder downstream phylogenetic analyses.

### The genus *Microniphargus* is more closely related to *Pseudoniphargus* than to *Niphargus*

Our COI and 28S phylogenetic trees show that *Microniphargus* is monophyletic and clearly distinct from *Niphargus* and *Pseudoniphargus*, thereby confirming its status as a separate genus established on morphological characters (Schellenberg 1934). The results of our analysis confirm the conclusions reached by Moškrič & Verovnik (2019) and Copilaş-Ciocianu et al. (2020) on the close affinity between *Microniphargus* and *Pseudoniphargus*, suggesting the inclusion of the genus *Microniphargus* within the family Pseudoniphargidae to avoid paraphyly of Niphargidae. Consequently, superfamilies Allocrangonyctoidea and Crangonyctoidea as proposed by Lowry & Myers (2012, 2017) turn out to be paraphyletic.

As mentioned in the introduction, a similarity between the two genera can be found in the shape of the telson (which is widely incised and carries one spine on each lobe), and also partly the shape of gnathopod 1. This shape of telson as well as the protrusion on the carpus of gnathopod 1 are found also in Bogidiellidae (another family placed in recent phylogenetic trees not far away from the clade Niphargidae+Pseudoniphargidae: Copilaş-Ciocianu et al., 2020) and may be simply symplesiomorphic, in which case the deeply incised, bilobated telson of *Niphargus* would represent an apomorphic character of this genus. However, the small size of *Microniphargus*, the reduced setation of mandibular palp and gnathopods, the lack of elongation of the third uropod in males, and the 1-articulated accessory flagellum of antennulae suggest a major role of paedomorphosis, making it difficult or impossible to correctly allocate this genus within current amphipod taxonomy and phylogeny based on morphological characters alone.

The inclusion of *Microniphargus* within Pseudoniphargidae requires an adjustment in the diagnosis of the family, recently revised by Lowry & Myers (2012), with minor changes as follows:

Body depigmented; eyes absent. Antenna 1 longer than antenna 2; accessory flagellum short, or minute, 1-2 articulated. Gnathopod 1 smaller (or weaker) than gnathopod 2; propodus with multiple groups of simple or bifid setae along palmar margin. Urosomites 1 to 3 free, without robust dorsal setae. Urosomite 1 without distoventral robust seta. Uropod 3 biramous; inner ramus minute; outer ramus article 2 absent. Telson notched, distal margin emarginate or nearly straight, with 1-3 robust spines on each lobe.

### *Microniphargus leruthi* comprises three cryptic lineages

Our COI phylogeny, ABGD’s initial partitioning of our comprehensive COI dataset and our haploweb analysis of 28S sequences of *Microniphargus* support the hypothesis that *Microniphargus leruthi* is composed of three distinct, putatively species-level lineages: clade A found in Ireland, clade B found both in England and in Belgium (with two COI sub-clades consistent with the geographic distance between these two locations), and clade C found so far only in England. Although ABGD’s recursive partitioning supports a four-species hypothesis, the p-distances between these three main lineages are all well above the 3% species-level threshold traditionally considered in barcoding studies (Hebert *et al.* 2003), whereas the average p-distance between the two COI sub-clades of B falls below this symbolic threshold. These arguments, together with the fact that all individuals of lineage B (and only these individuals) display double peaks in their COI chromatograms, lead us to consider tentatively the two sub-clades of lineage C as conspecific and therefore to distinguish only three putative species-level lineages within *M. leruthi*.

Although lineage A (found only in Ireland to date) appears geographically separated from the other two, lineages B and C occur in sympatry in at least one location (Swildon’s Hole in Somerset), bringing further support to the hypothesis that these two lineages are distinct species. The phylogenetic analysis based on COI marker strongly suggest an origin of the genus *Microniphargus* in England, with subsequent dispersals to Ireland and to Belgium. The fact that lineage B still occurs on both sides of the English Channel is not overly surprising since the land connection between England and continental Europe was only severed about 8,000 years ago (Waller & Long 2003).

The hypothesis that the three *Microniphargus leruthi* lineages identified here represent distinct cryptic (or pseudo-cryptic) species will need to be tested further. Doing so will require further collecting and sequencing, as well as detailed morphological analyses using microscopy techniques appropriate for such small specimens.

## Acknowledgements

Thanks to John Boulton and Camille Ek for helping with fieldwork and collecting.

## Supplementary material

**Figure S1:**
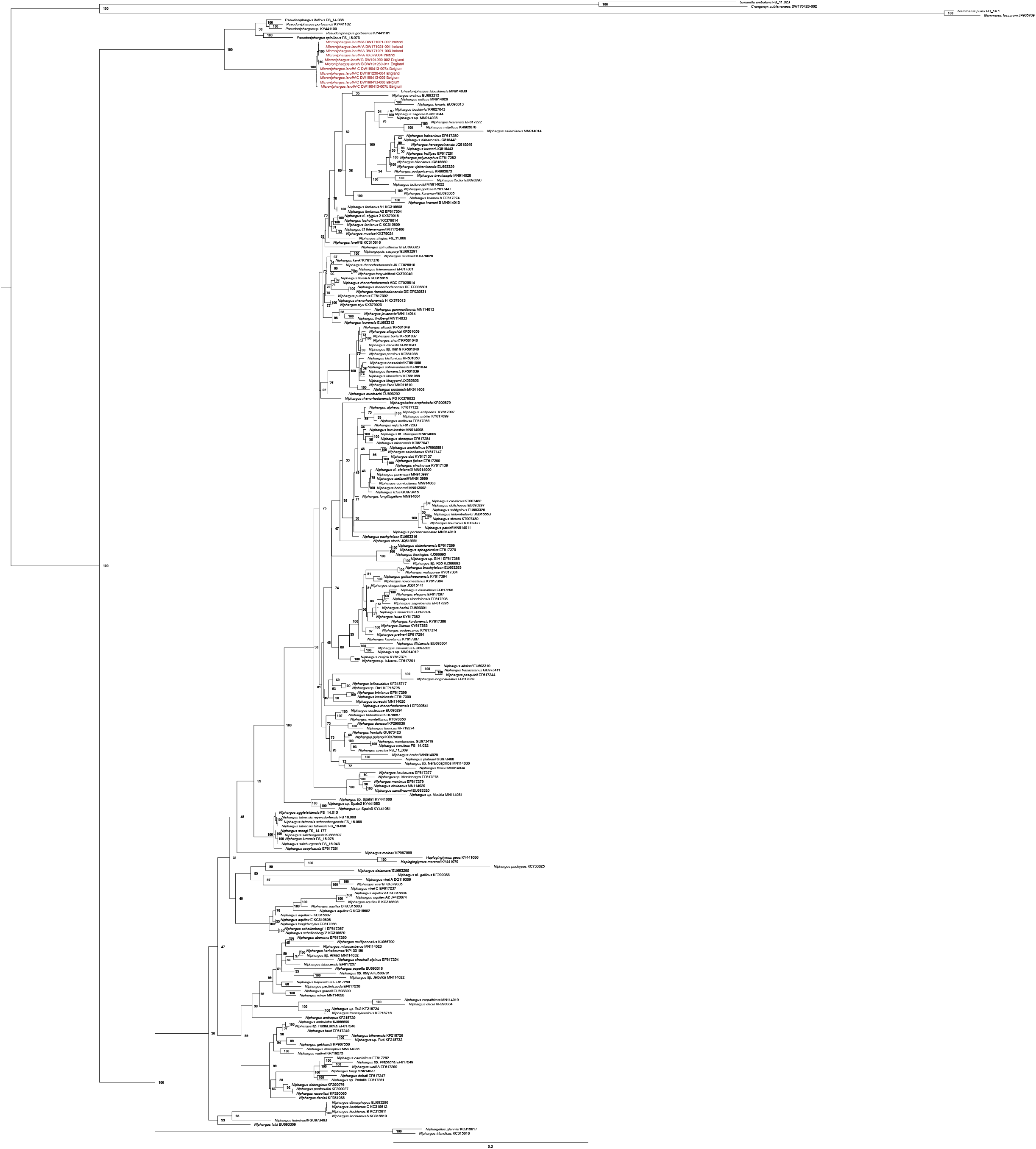
Complete, uncondensed version of the 28S maximum-likelihood phylogeny in Figure 3

**Figure S2:**
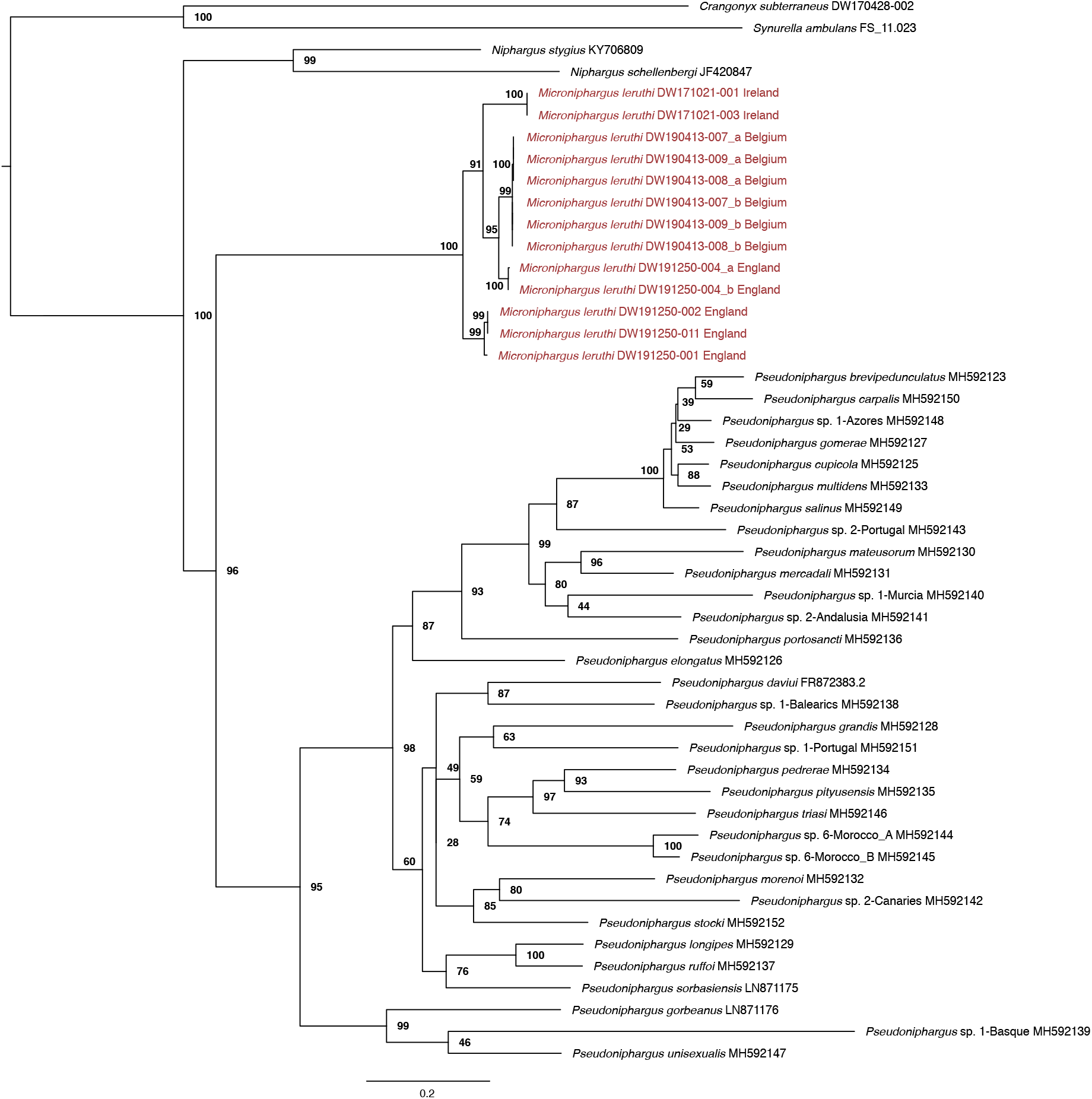
Complete, uncondensed version of the COI maximum-likelihood phylogeny in Figure 4

**Table S1:**
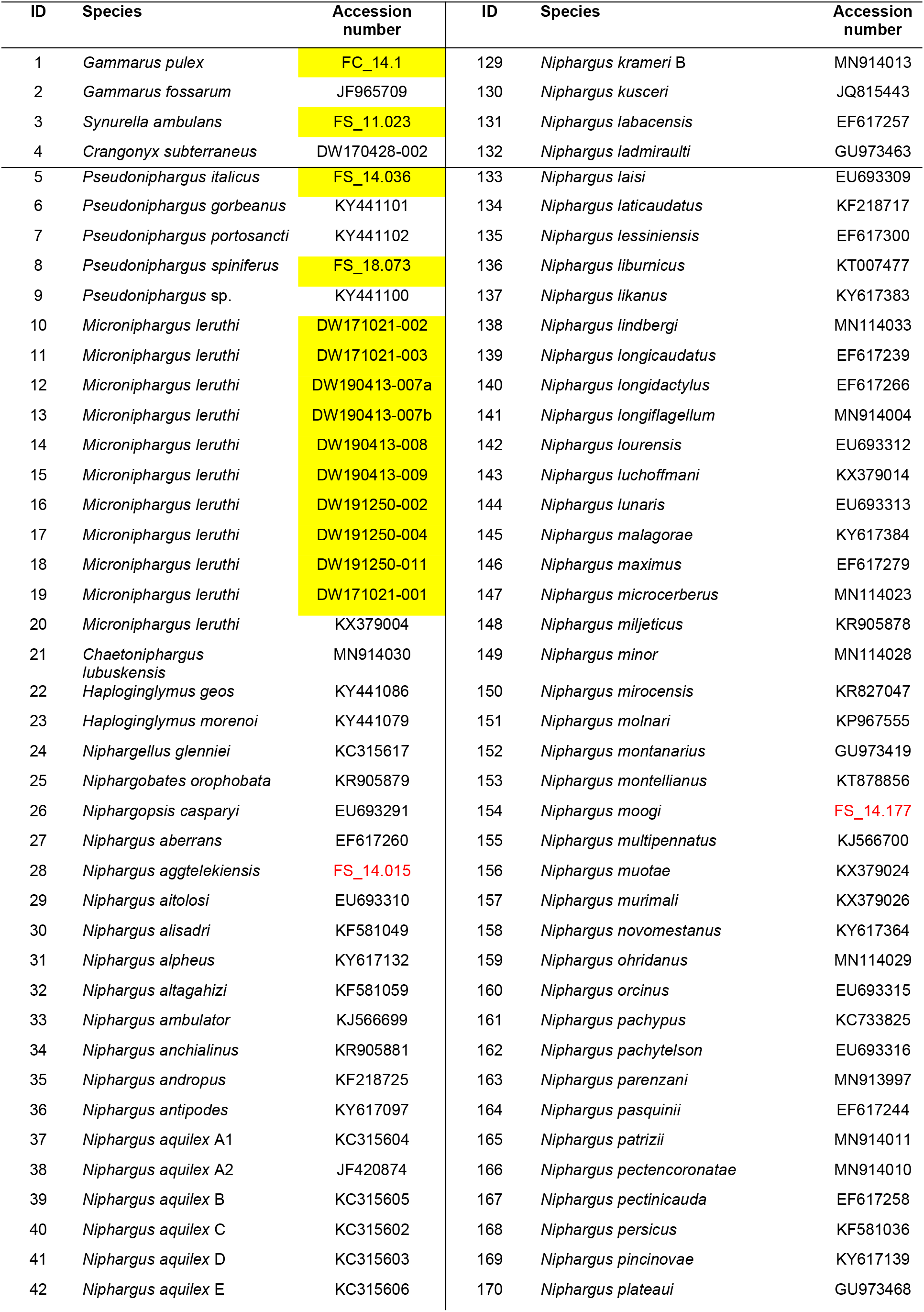
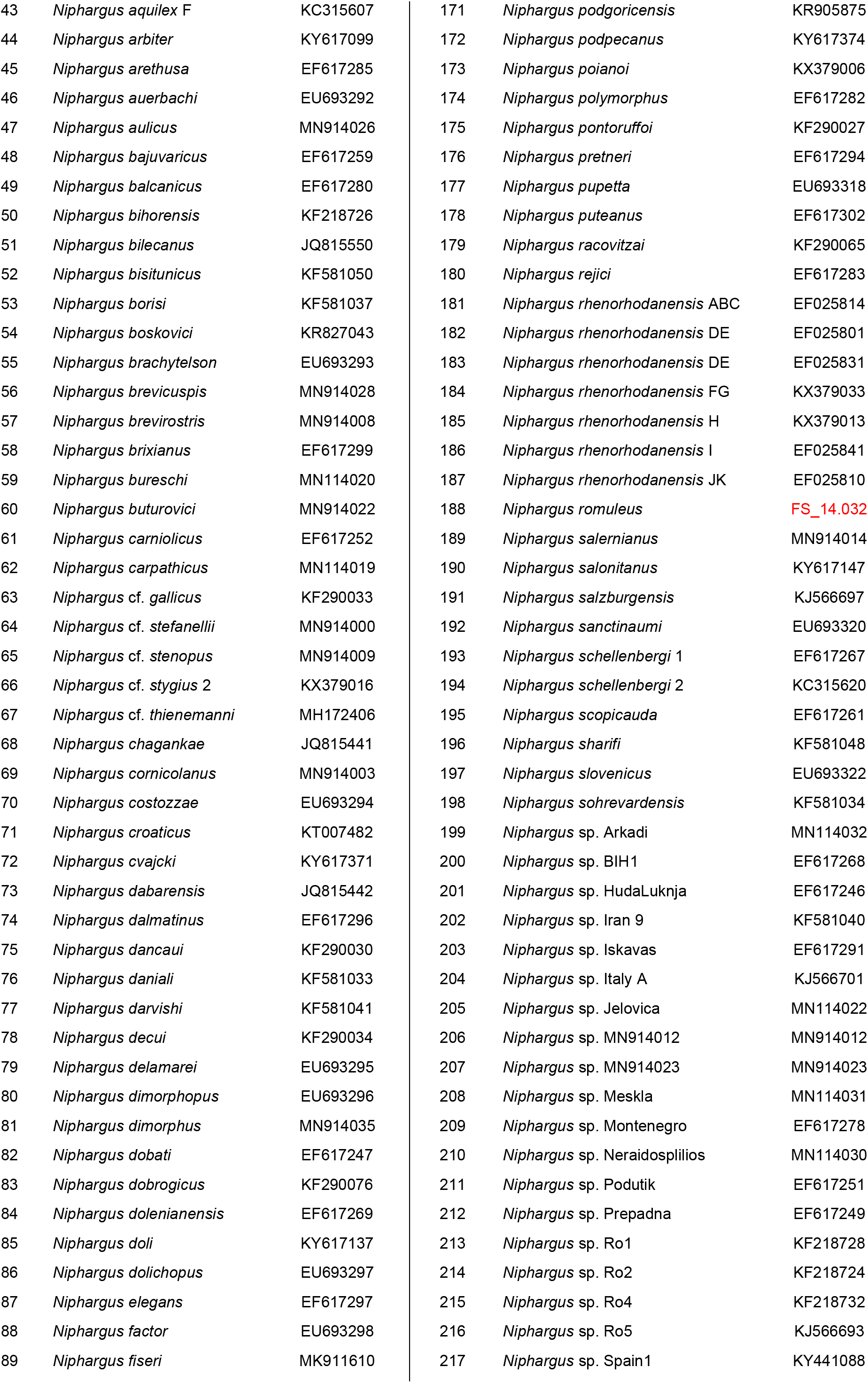
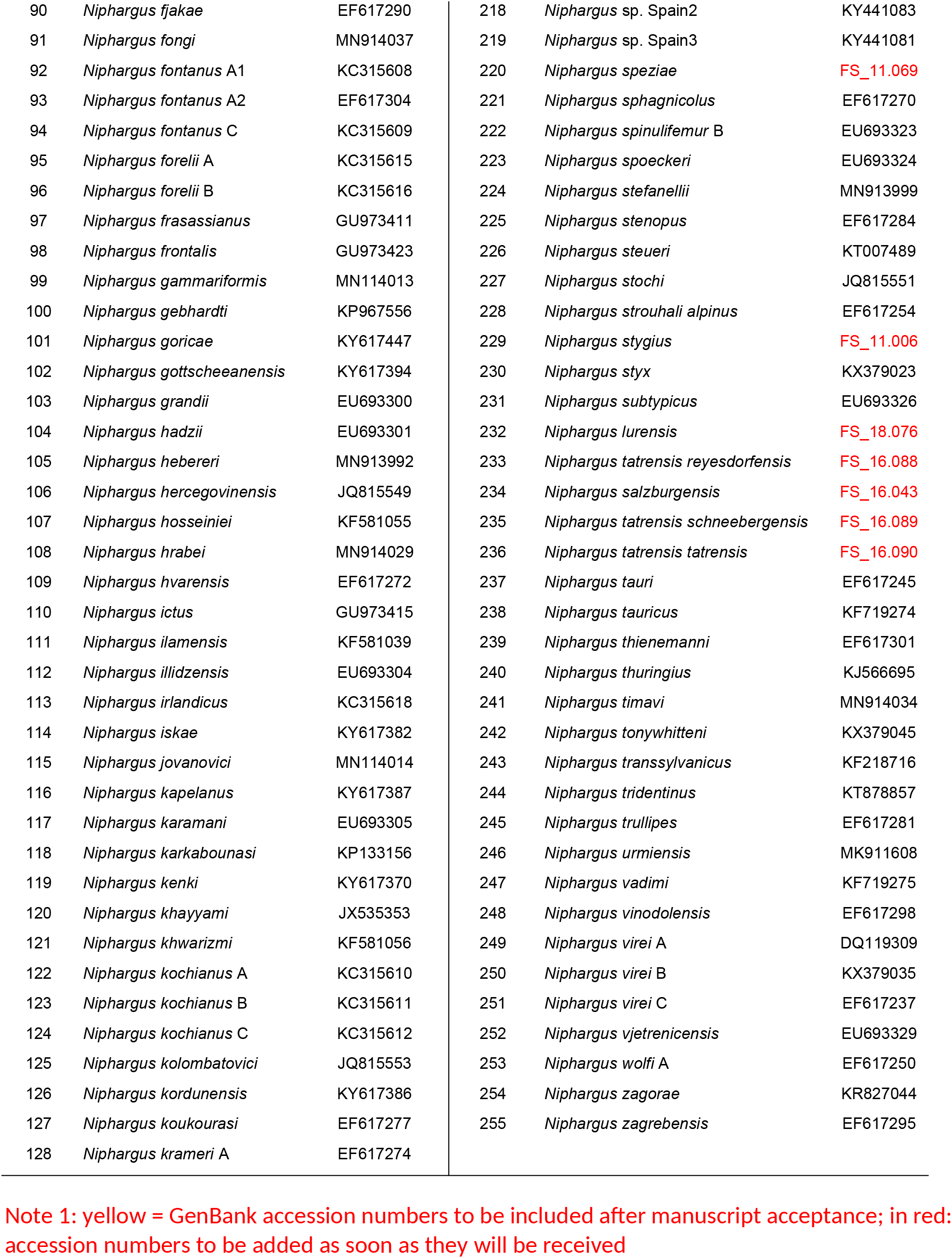
List of all the sequences included in the 28S phylogeny, including species names and GenBank accession numbers

**Table S2:**
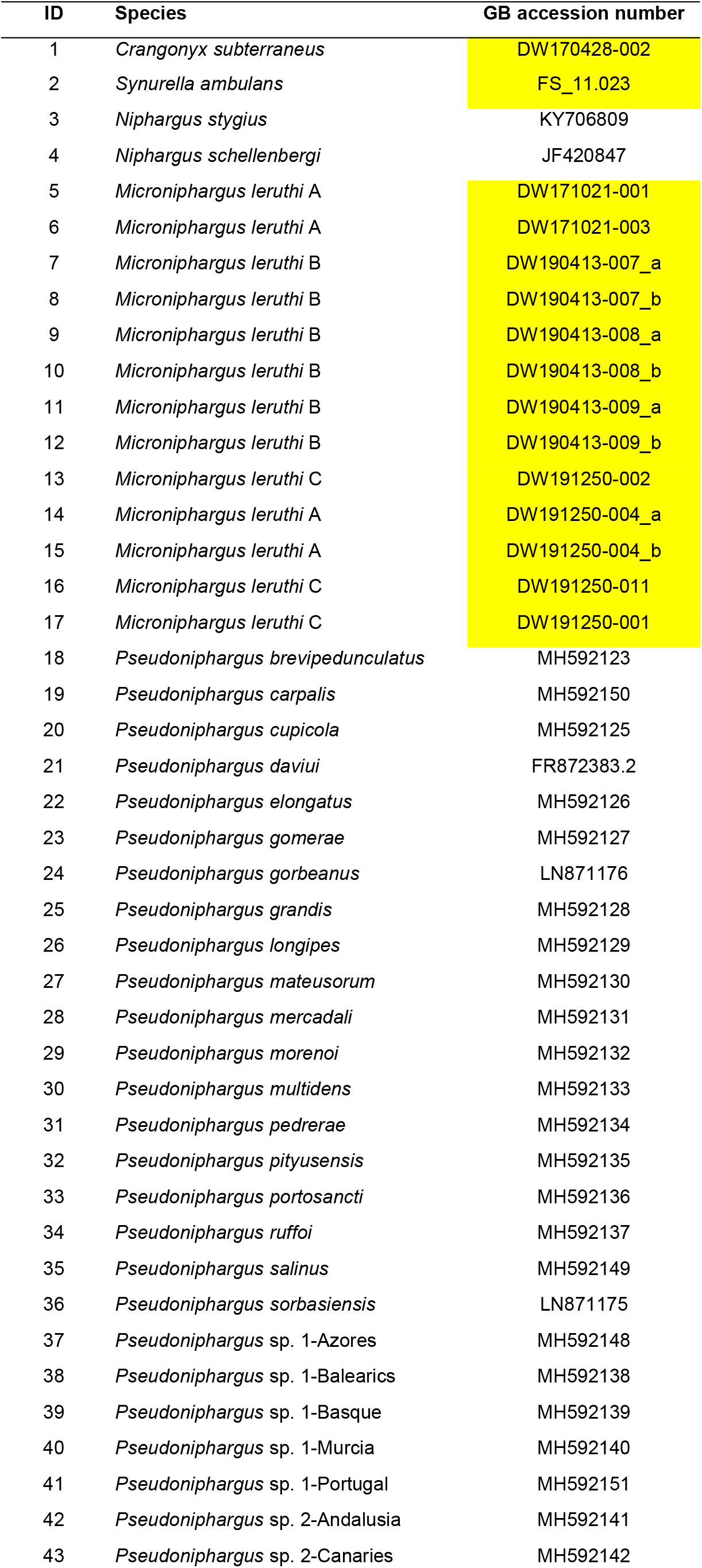
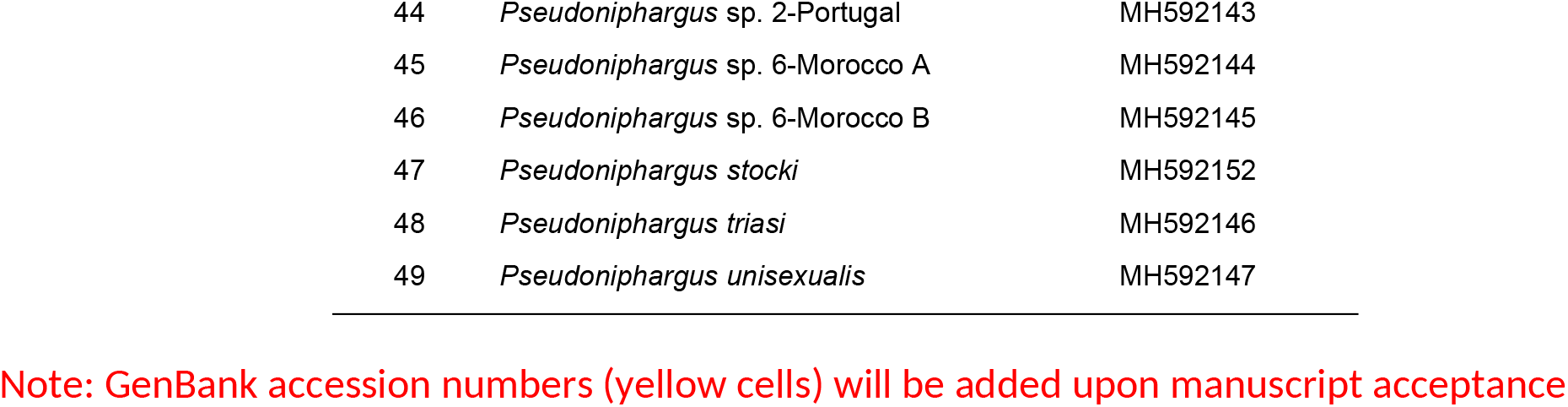
List of all the sequences included in the COI phylogeny, including species names and GenBank accession numbers

